# A quinoa-associated *Pantoea* isolate displays salinity-responsive auxin production and promotes plant growth under salt stress

**DOI:** 10.64898/2026.07.02.736047

**Authors:** Yoshinori Murata, Takeshi Kashiwa, Hatairat Dangjarean, Yasufumi Kobayashi, Yasunari Fujita

**Author notes:** Corresponding author. Yasunari Fujita.

## Abstract

Plant-associated bacteria can promote plant growth under saline conditions, but salinity-dependent changes in bacterial physiological traits remain insufficiently understood. Here, we isolated bacteria from seedlings of quinoa (*Chenopodium quinoa* Willd.) lines maintained under laboratory propagation for more than 30 years and evaluated their activity under saline conditions. A quinoa-associated *Pantoea* isolate, strain 6PN, promoted primary root elongation and whole-plant dry weight of *Arabidopsis thaliana* under salt stress, whereas no significant effect was observed under non-saline conditions. Comparative analyses with reference *Pantoea agglomerans* strains showed that strain 6PN exhibited salinity-responsive indole-3-acetic acid (IAA) production. Genome analysis identified a putative *ipdC* gene and additional genes related to stress responses, nutrient acquisition, polysaccharide biosynthesis and export, flagellar biosynthesis, and chemotaxis. Phylogenomic analysis indicated that strain 6PN was genomically distinct from representative *Pantoea* species examined here. In an *Arabidopsis* trench-plate assay, GFP-labeled strain 6PN was recovered from spatially separated plant tissues at higher levels than a GFP-labeled reference strain under saline conditions. These results identify strain 6PN as a quinoa-associated *Pantoea* isolate with salinity-responsive IAA production and plant growth-promoting activity under defined salt-stress conditions.

## Introduction

Salinity stress is one of the major environmental factors limiting agricultural productivity worldwide and is expected to become increasingly severe under climate change and soil degradation^1,2^. Plants growing under saline environments exhibit complex physiological and developmental adaptations involving ion homeostasis, osmotic adjustment, hormonal regulation, and reactive oxygen species management^1,3,4^. In recent years, plant-associated microorganisms, particularly plant growth-promoting rhizobacteria, have been increasingly recognized as important contributors to plant performance under stressful environmental conditions, including salinity and drought stress^5,6^.

A wide variety of plant-associated bacteria have been reported to promote plant growth through mechanisms such as phytohormone production, phosphorus solubilization, siderophore production, and stress-related signaling^5,7,8^. Among these mechanisms, bacterial production of indole-3-acetic acid (IAA) has attracted particular attention because of its potential effects on root development and stress-associated growth responses^9,10^. In plants, the indole-3-pyruvic acid (IPyA) pathway mediated by TAA1/TAR and YUCCA enzymes represents a major endogenous route of IAA biosynthesis involved in growth and developmental regulation^11^. In contrast, bacteria utilize multiple IAA biosynthesis pathways, including IPyA-, indole-3-acetamide (IAM)-, tryptamine (TAM)-, and indole-3-acetonitrile (IAN)-associated pathways, depending on the bacterial species and environmental conditions^12^. However, much of the previous work has primarily focused on whether bacterial strains produce IAA, whereas comparatively little is known about how environmental stress conditions dynamically influence bacterial auxin biosynthesis pathways and associated plant growth-promoting activity^12–14^.

Quinoa (*Chenopodium quinoa* Willd.) is an Andean orphan crop that has attracted increasing attention as a climate-resilient crop and is widely recognized for its exceptional tolerance to salinity, drought, and other harsh environmental conditions^15^. Previous studies have demonstrated that quinoa possesses strong intrinsic salinity tolerance, including tolerance to seawater-level salinity conditions^16,17^. Notably, quinoa lines maintained under laboratory propagation conditions for more than 30 years still retain robust salinity tolerance^17,18^, indicating that quinoa possesses stable intrinsic adaptive mechanisms. At the same time, plants growing under stressful environments often harbor microorganisms with stress-associated physiological functions. Recent studies have suggested that microbiota associated with stress-adapted plants can influence plant growth under stressful environments^5,19–21^. However, little is known about the physiological traits of bacteria isolated from quinoa plants derived from long-term laboratory-propagated lines, or how environmental salinity affects bacterial traits associated with plant growth promotion.

Here, we isolated plant-associated bacteria from quinoa lines maintained under long-term laboratory propagation and evaluated their plant growth-promoting activity under saline conditions. We identified a quinoa-associated *Pantoea* isolate, strain 6PN, that promoted plant growth under salt stress and exhibited salinity-responsive IAA production. We further performed comparative physiological assays, genome analysis, phylogenomic analysis, and an *Arabidopsis* trench-plate assay to describe bacterial traits associated with salinity-responsive IAA production, plant growth-promoting activity, and recovery from spatially separated plant tissues under defined saline conditions.

## Materials and methods

### Isolation of microorganisms from quinoa plants

Quinoa was cultivated as described previously^17,22,23^, with minor modifications. Seeds of the quinoa Kd inbred line were used in this study. The Kd line was established at JIRCAS as the inbred accession Kyoto-d from quinoa seeds that had been propagated for more than 20 years in a temperature-controlled plant growth room at Kyoto University in the absence of other quinoa accessions^18^. At JIRCAS, progeny seeds derived from a single plant were further propagated under controlled conditions using non-woven pollination bags to prevent cross-pollination^18^. Seeds of the Kd line used here had been maintained under laboratory propagation conditions for more than 30 years in total. Seeds were surface-sterilized with 70% ethanol for 5–10 min, followed by immersion in 1.0% sodium hypochlorite solution for 5–10 min, and subsequently rinsed four times with sterile ultrapure water. The sterilized seeds were placed on half-strength Murashige and Skoog (0.5× MS) agar medium supplemented with 2% (w/v) sucrose and incubated at 25°C until germination. Six days after germination, quinoa seedlings were transferred to cell trays containing a sterilized substrate composed of vermiculite and potting mix for professionals (Innovex, Tokyo, Japan) at a ratio of 9:1 (v/v). The substrate had been autoclaved and dried prior to use. Seedlings were watered for 2 days and subsequently cultivated under salt stress conditions (300 mM NaCl) at 25°C for 7 days. After salt treatment, seedlings were surface-sterilized by washing three times with sterile ultrapure water, followed by immersion in 70% ethanol for 1 min, 1% sodium hypochlorite for 1 min, and two additional washes with sterile ultrapure water. To confirm the effectiveness of this final surface-sterilization step, the final rinse water was spread onto Reasoner’s 2A (R2A) agar plates and incubated at 25°C for 3 days; no colony formation was observed. Residual moisture was removed using sterile paper towels. The sterilized tissues were homogenized with sterile zirconia beads using a Shake Master homogenizer (BMS-A20TP, BioMedical Science, Tokyo, Japan). The resulting homogenates were spread onto tryptic soy agar (TSA) and R2A agar plates and incubated at 25°C for 4 days. Distinct bacterial colonies that appeared on the plates were isolated and purified. Based on preliminary tests, we selected bacterial strain 6PN as a candidate strain that promotes plant growth under salt stress.

### Identification and phylogenetic analysis of strain 6PN

The isolated bacterium, strain 6PN, was initially identified by PCR amplification and sequencing of the 16S rRNA gene. Genomic DNA was extracted using the Wizard Genomic DNA Purification Kit (Promega, WI, USA). PCR amplification was performed in a 50 μL reaction mixture containing 25 μL of 2× MightyAmp Buffer Ver.3, 2 μL of template DNA (0.1–0.2 μg), 1 μL of MightyAmp DNA Polymerase Ver.3, 1 μL each of the forward primer 27F (5′-AGAGTTTGATCCTGGCTCAG-3′) and reverse primer 1492R (5′-GGTTACCTTGTTACGACTT-3′), and 21 μL of sterile ultrapure water.

PCR amplification was carried out under the following conditions: initial denaturation at 98°C for 2 min; 35 cycles of denaturation at 98°C for 10 s, annealing at 58°C for 15 s, and extension at 68°C for 1.5 min; followed by a final extension at 68°C for 5 min. The obtained 16S rRNA gene sequence was compared with sequences in the NCBI nucleotide database using BLAST analysis. Based on sequence similarity, strain 6PN was identified as a member of the genus *Pantoea*.

For 16S rRNA gene-based phylogenetic analysis, the 16S rRNA gene sequence of strain 6PN was retrieved from the genome assembly generated in this study and compared with 16S rRNA gene sequences of representative *Pantoea* strains available in the NCBI database (https://www.ncbi.nlm.nih.gov/, accessed on June 3–10, 2026): *P. agglomerans* FDAARGOS 1447 (= DSM 3493 = NBRC 102470^T^; accession no. CP077366.1), *P. allii* LMG 24248ᵀ (CP193910.1), *P. eucalypti* LMG 24197ᵀ (CP045720.1), and *P. vagans* LMG 24199ᵀ (CP038853.1). *Erwinia amylovora* ATCC 15580ᵀ (CP066796.1) was used as the outgroup. Sequence alignment was performed using MAFFT v7.490^24^ implemented in Geneious Prime 2026.0.2 with the L-INS-i algorithm. The alignment was trimmed at one end to remove a poorly aligned terminal region and to standardize the sequence boundaries. A maximum-likelihood phylogenetic tree was constructed using IQ-TREE v3.1.2^25^ with the HKY substitution model selected by the software. Branch support was assessed using 1,000 bootstrap replicates. The resulting tree was visualized using FigTree v1.4.4 (https://github.com/rambaut/figtree, accessed on June 2, 2026).

### Bacterial inoculation onto plants

Seeds of *Arabidopsis thaliana* CS60000 were surface-sterilized according to Nagatoshi et al. (2023)^26^ and sown on germination medium consisting of 0.5× MS medium. Four days after germination, seedlings were transferred to plant growth assay medium consisting of 0.5× MS medium without sucrose and containing 3% agar, or to salinity stress medium consisting of the same medium supplemented with 100–150 mM NaCl. Seedlings were cultivated at 22°C under long-day conditions (16 h light/8 h dark). Strain 6PN was cultured on TSA plates for 2–3 days. A single colony was inoculated into tryptic soy broth and incubated for 24 h. The liquid culture was subsequently transferred into fresh tryptic soy broth and further incubated at 25°C. After 6 h of cultivation, bacterial cells were collected by centrifugation and washed twice with 10 mM MgCl₂ to remove residual medium components. The resulting pellet was resuspended in 10 mM MgCl₂, and the optical density at 600 nm (OD600) was adjusted to 0.1. The bacterial suspension was diluted with 10 mM MgCl₂ to the indicated concentrations, and 5 μL of the diluted suspension was inoculated onto the root region of *A. thaliana* seedlings 2 d after transfer to treatment medium. For comparison with strain 6PN in whole-plant dry weight assays, *Pantoea agglomerans* NBRC 102470 was prepared and inoculated in the same manner. All plant growth assays were independently repeated at least three times. Data shown in Fig. 2 are representative of three independent experiments that yielded consistent results.

### Evaluation of plant growth-promoting traits

To compare the plant growth-promoting traits of strain 6PN with those of closely related reference strains, *Pantoea agglomerans* NBRC 102470 and NBRC 12686 were obtained from the Biological Resource Center, National Institute of Technology and Evaluation (NBRC), Japan. These strains were selected as reference strains because they showed high 16S rRNA gene sequence similarity to strain 6PN and were available from a public culture collection. Both strains were used in the following assays alongside strain 6PN. Plant growth-promoting traits were evaluated under non-saline and saline conditions. Phosphate solubilization, CMC degradation, and siderophore production assays were performed in the absence or presence of 100 mM NaCl. IAA production was evaluated in one-tenth-strength TSB supplemented with 1 mM tryptophan, with or without 100 mM NaCl, as described below.

### Phosphate solubilization assay

Phosphate solubilization activity was evaluated using phosphate agar medium consisting of 10.0 g glucose, 0.5 g (NH₄)₂SO₄, 0.5 g yeast extract, 0.3 g NaCl, 0.3 g KCl, 0.03 g FeSO₄·7H₂O, 0.3 g MgSO₄·7H₂O, 0.03 g MnSO₄·4H₂O, 5.0 g Ca₃(PO₄)₂, and 15.0 g agar per liter of ultrapure water (pH 7.0–7.5)^27^. Bacterial suspensions of strain 6PN and reference strains were prepared in 10 mM MgCl₂ and adjusted to OD600 = 0.05. Bacterial suspensions were inoculated onto phosphate agar plates at 3–4 μL per spot. After incubation for 7 days, phosphate solubilization activity was evaluated by measuring the halo diameter surrounding bacterial colonies.

### Carboxymethyl cellulose degradation assay

Carboxymethyl cellulose (CMC) degradation activity was evaluated using CMC-Na agar medium consisting of 2.0 g KH₂PO₄, 1.4 g (NH₄)₂SO₄, 0.3 g MgSO₄·7H₂O, 0.3 g CaCl₂, 0.4 g yeast extract, 0.005 g FeSO₄·7H₂O, 0.0016 g MnSO₄, 0.0017 g ZnCl₂, 0.002 g CoCl₂, 5.0 g CMC-Na, and 15.0 g agar per liter of ultrapure water (pH 5.0)^28^. Bacterial suspensions were prepared in 10 mM MgCl₂ and adjusted to OD600 = 0.05. Bacterial suspensions were inoculated onto CMC agar plates at 3–4 μL per spot and incubated for 7 days. Plates were subsequently stained with 5% Ponceau S solution, and the halo diameter surrounding colonies was measured as an indicator of CMC degradation activity.

### Siderophore production assay

Siderophore production was evaluated using chrome azurol S (CAS) agar according to the method of Louden et al.^29^ with minor modifications. For preparation of the CAS blue dye solution, 0.06 g chrome azurol S (CAS; MP Biomedicals, CA, USA) was dissolved in 50 mL ultrapure water. Separately, 0.0027 g FeCl₃·6H₂O was dissolved in 10 mL of 10 mM HCl, and 0.073 g hexadecyltrimethylammonium bromide (HDTMA) was dissolved in 40 mL ultrapure water. The CAS solution was mixed with 9 mL of the FeCl₃ solution, followed by addition of the HDTMA solution. For preparation of the agar base, MM9 salt stock was prepared by dissolving 15 g KH₂PO₄, 25 g NaCl, and 50 g NH₄Cl in 500 mL ultrapure water. A 20% glucose stock solution was prepared separately. Casamino acids (3 g) were dissolved in 27 mL ultrapure water and sterilized by autoclaving. Subsequently, 100 mL MM9 salt stock was added to 750 mL ultrapure water, followed by dissolution of 32.24 g PIPES buffer. The pH was adjusted to 6.8, and 15 g Bacto agar was added before autoclaving. After cooling to approximately 50°C, sterile casamino acid solution (30 mL), sterile 20% glucose solution (10 mL), and sterile CAS blue dye solution (100 mL) were added with gentle mixing. Plates were poured aseptically and allowed to solidify. For siderophore assays, 3–4 μL of bacterial suspension was inoculated onto CAS agar plates. Colony diameter and surrounding orange halo diameter were measured daily from 1 to 4 days after inoculation.

### Indole-3-acetic acid (IAA) production assay

Strains 6PN, NBRC 102470, and NBRC 12686 were cultured overnight in tryptic soy broth (TSB) at 25°C. The overnight cultures were subsequently reinoculated into fresh TSB and incubated for an additional 6 h at 25°C. Bacterial cells were collected and washed three times with one-tenth-strength TSB (1/10 TSB) prepared by diluting TSB 10-fold with sterile ultrapure water. The washed cells were resuspended in 1/10 TSB and inoculated into the following two culture conditions: (i) 1/10 TSB supplemented with 1 mM tryptophan and (ii) 1/10 TSB supplemented with 1 mM tryptophan and 100 mM NaCl. The final cell density was adjusted to OD600 = 0.05, and cultures were incubated at 25°C with shaking at 150 rpm. For the tryptophan dose-response assay, strain 6PN was cultured in 1/10 TSB supplemented with the indicated concentrations of tryptophan, and IAA accumulation was quantified by HPLC as described below.

For IAA quantification, 0.5 mL culture supernatant was collected at 24, 48, and 72 h after inoculation and centrifuged at 15,000 × g for 10 min at 4°C. The resulting supernatants were directly subjected to high-performance liquid chromatography (HPLC) analysis. The HPLC system consisted of a binary pump (PU-4185 Binary; Jasco, Tokyo, Japan), a fluorescence detector (FP-4025; Jasco), an autosampler (AS-4150; Jasco), and a column oven (CO-4060; Jasco). IAA quantification was performed based on a modified method of Szkop and Bielawski (2013)^30^. Separation was carried out using an Inertsil C8 column (5 μm, 4.6 mm i.d. × 150 mm; GL Sciences, Tokyo, Japan) equipped with a guard cartridge column E (4.0 mm × 10 mm; GL Sciences) at 40°C. Solvent A consisted of 2.5:97.5% (v/v) acetic acid:H₂O (pH 3.8 adjusted with 1 M KOH), and solvent B consisted of 80:20% (v/v) acetonitrile:H₂O. Gradient elution was performed using solvent A/solvent B at 80:20 from 0 to 20 min, followed by 50:50 from 21 to 35 min, 0:100 from 36 to 46 min, and re-equilibration at 80:20 from 47 to 60 min. The flow rate was maintained at 1.0 mL min⁻¹, and the maximum run time was 60 min. The injection volume was 40 μL. Fluorescence detection was performed with excitation and emission wavelengths set at 280 and 350 nm, respectively. Standard compounds used for analysis included indole-3-acetic acid (IAA), indole-3-acetonitrile (IAN), indole-3-acetamide (IAM), tryptamine (TAM), and indole-3-ethanol (TOL) (Sigma-Aldrich, NJ, USA). Quantification of IAA was performed using a standard calibration curve ranging from 0.1 to 100 nmol mL⁻¹. Data acquisition and quantification were conducted using ChromNAV software version 2.0 (Jasco, Tokyo, Japan).

### Genome sequencing, phylogenomic analysis, and ANI analysis of strain 6PN

Genomic DNA of strain 6PN was extracted from a 19-h bacterial culture using the NucleoBond HMW DNA kit (Macherey-Nagel, Düren, Germany). Purified DNA was quantified using the Qubit 1× dsDNA HS Assay Kit (Thermo Fisher Scientific, Waltham, MA, USA). Sequencing libraries were prepared using the Ligation Sequencing Kit (SQK-LSK109, Oxford Nanopore Technologies, Oxford, UK) according to the manufacturer’s instructions. Genome sequencing was performed using three Flongle Flow Cells (FLO-FLG001, Oxford Nanopore Technologies) for a total run time of 70 h. Base calling was conducted using Guppy v6.0.1 (Oxford Nanopore Technologies) with default parameters. Genome assembly was performed using Flye v2.9 with the options “--nano-raw” and “--iterations 1”; the estimated genome size was set to 4.5 Mb during assembly. Contigs were polished once using Medaka v1.5.0 (Oxford Nanopore Technologies). Genome quality assessment, gene prediction, and functional annotation were conducted using the DFAST pipeline (https://dfast.ddbj.nig.ac.jp/, accessed on January 31, 2022). Genome maps were visualized using circos v0.69-8^31^.

The genome sequence of strain 6PN and RefSeq chromosomal sequences retrieved from the NCBI database were used for genome-based phylogenetic analysis. The RefSeq chromosomal sequences used for extracting housekeeping genes included *Pantoea allii* Eh250 (accession no. NZ_CP125958.1), *Pantoea agglomerans* FDAARGOS 1447 (= DSM 3493 = NBRC 102470; NZ_CP077366.1), *Pantoea eucalypti* LMG 24197 (NZ_CP045720.1), *Pantoea vagans* LMG 24199 (NZ_CP038853.1), *Pantoea ananatis* PA13 (NC_017554.1), and *Erwinia amylovora* EaSmR (NZ_CP171267.1). Although *P. agglomerans* NBRC 102470 was used as a reference strain in physiological and plant-association assays, the genome assembly deposited for NBRC 102470 was not used for genome-based phylogenetic analysis because it was highly fragmented. Instead, the RefSeq chromosomal sequence of *P. agglomerans* FDAARGOS 1447, corresponding to DSM 3493, the type strain of *P. agglomerans*, was used. Coding sequences of six housekeeping genes (*rpoD*, *rpoB*, *recA*, *dnaN*, *dnaA*, and *atpD*) were extracted from these chromosomal sequences and used to construct a multilocus phylogenetic tree. The concatenated alignment was generated using MAFFT v7.490^24^ implemented in Geneious Prime 2026.0.2 with the L-INS-i algorithm. Unreliably aligned regions were removed using Gblocks v1.0^32,33^. A maximum-likelihood phylogenetic tree was constructed using IQ-TREE v3.1.2 (Wong et al., 2026) ^25^ with the TIM2+F+I+G4 model selected by IQ-TREE. Branch support was assessed using 1,000 bootstrap replicates. The resulting tree was visualized using FigTree v1.4.4 (https://github.com/rambaut/figtree, accessed on June 2, 2026). Average nucleotide identity (ANI) was calculated using the genome assembly of strain 6PN and whole-genome assemblies of representative *Pantoea* strains. The strains included in the ANI matrix were selected based on their close phylogenetic proximity to strain 6PN and the availability of whole-genome assemblies, and included *P. vagans* LMG 24199 (accession no. GCA_004792415.1), *P. agglomerans* FDAARGOS 1447 (= DSM 3493 = NBRC 102470; GCA_019048385.1), *P. eucalypti* LMG 24197 (GCA_009646115.1), and *P. alhagi* LTYR-11Z (GCA_002101395.1). ANI values were calculated using OrthoANI v0.5.0^34^.

### Evaluation of bacterial association with plant tissues under saline conditions

The plasmid pBBR1-MCS5-GFP was kindly provided by Dr. Jennifer L. Morrell-Falvey (Biosciences Division, Oak Ridge National Laboratory, Oak Ridge, TN, USA)^35^. Strains 6PN and NBRC 102470 were transformed with pBBR1-MCS5 carrying a constitutively expressed green fluorescent protein (GFP) gene, generating the GFP-labeled strains 6PN-GFP and NBRC 102470-GFP. For electroporation, 20 ng of plasmid DNA in 2.0 μL was mixed with 60 μL of competent *Pantoea* cells. Electroporation was performed in a 0.2-cm-gap cuvette at 20–30 kV cm⁻¹, with a capacitance of 23 μF and a resistance of 200 Ω. GFP-labeled strains were maintained on medium supplemented with 200 μg mL⁻¹ gentamicin.

Seeds of *Arabidopsis thaliana* CS60000 were surface-sterilized according to Nagatoshi et al. (2023)^26^ and sown on germination medium consisting of 0.5× MS medium supplemented with 3% sucrose and 3% gellan gum. Five days after germination, seedlings were transferred onto trench plates containing 0.5× MS medium without sucrose and supplemented with 3% agar and 120 mM NaCl. Each plate contained a single trench positioned approximately one-third from the upper edge of the plate, physically separating the leaf and root regions. This system was used to evaluate bacterial association with spatially separated plant tissues under saline conditions. After transplantation, seedlings were cultivated at 22°C under long-day conditions (16 h light/8 h dark). GFP-labeled bacterial strains were cultured overnight in TSB containing 200 μg mL⁻¹ gentamicin. Bacterial cultures were centrifuged, and the resulting pellets were washed twice with 10 mM MgCl₂ to remove residual medium components. Cell suspensions were adjusted to OD600 = 0.025, and 3 μL of the suspension was inoculated onto either the leaf or root region of *A. thaliana* seedlings on trench plates. After 5–7 days of incubation, GFP fluorescence on plant tissues was observed using a fluorescence microscope (BZ-X810, KEYENCE, Osaka, Japan). For quantification of bacterial association, leaf or root tissues were excised, weighed, and homogenized with sterilized zirconia beads using a Shake Master homogenizer (BMS-A20TP, BioMedical Science, Tokyo, Japan). Homogenates were serially diluted with 10 mM MgCl₂ and spread onto TSA plates supplemented with 200 μg mL⁻¹ gentamicin. After 2 days of incubation, GFP-labeled colony-forming units (CFUs) were counted using a colony counter (aCOLyte 3 HD, Synbiosis, Cambridge, UK). Bacterial abundance was normalized to tissue fresh weight and expressed as CFUs per mg fresh weight.

### Statistical analysis

Data are presented as the mean ± standard deviation. The number of biological replicates for each experiment is indicated in the corresponding figure legends. Quantitative datasets were subjected to one-way analysis of variance (ANOVA) to detect overall differences among treatment groups, followed by Dunnett’s multiple comparisons test to compare each treatment with the control, Tukey’s HSD test to determine the level of significance among groups, or Student’s t-test to determine differences between two groups. All statistical analyses were performed using R (version 4.5.3) in RStudio (version 2026.01.2+418), and graphs were generated with the ggplot2 package^36^.

## Results

### A quinoa-associated *Pantoea* isolate promotes plant growth under salt stress

Preliminary assays using quinoa seedlings on salt-stress agar medium did not provide a suitable quantitative readout for bacterial growth-promoting activity. Quinoa seedlings showed strong intrinsic salinity tolerance and rapid root elongation on agar medium, making it difficult to define a salt-stress condition that reproducibly limited growth while still allowing sensitive detection of bacterial growth promotion. We therefore used *Arabidopsis thaliana* as a model plant system that enabled quantitative evaluation of bacterial growth-promoting activity under defined saline conditions. Among the bacterial isolates obtained from surface-sterilized young seedlings of the Kd quinoa line, one strain, designated 6PN, was initially identified as a member of the genus *Pantoea* by 16S rRNA gene sequencing. A maximum-likelihood phylogenetic tree based on 16S rRNA gene sequences further supported the placement of strain 6PN within the genus *Pantoea* (Fig. 1).

**Fig. 1.**
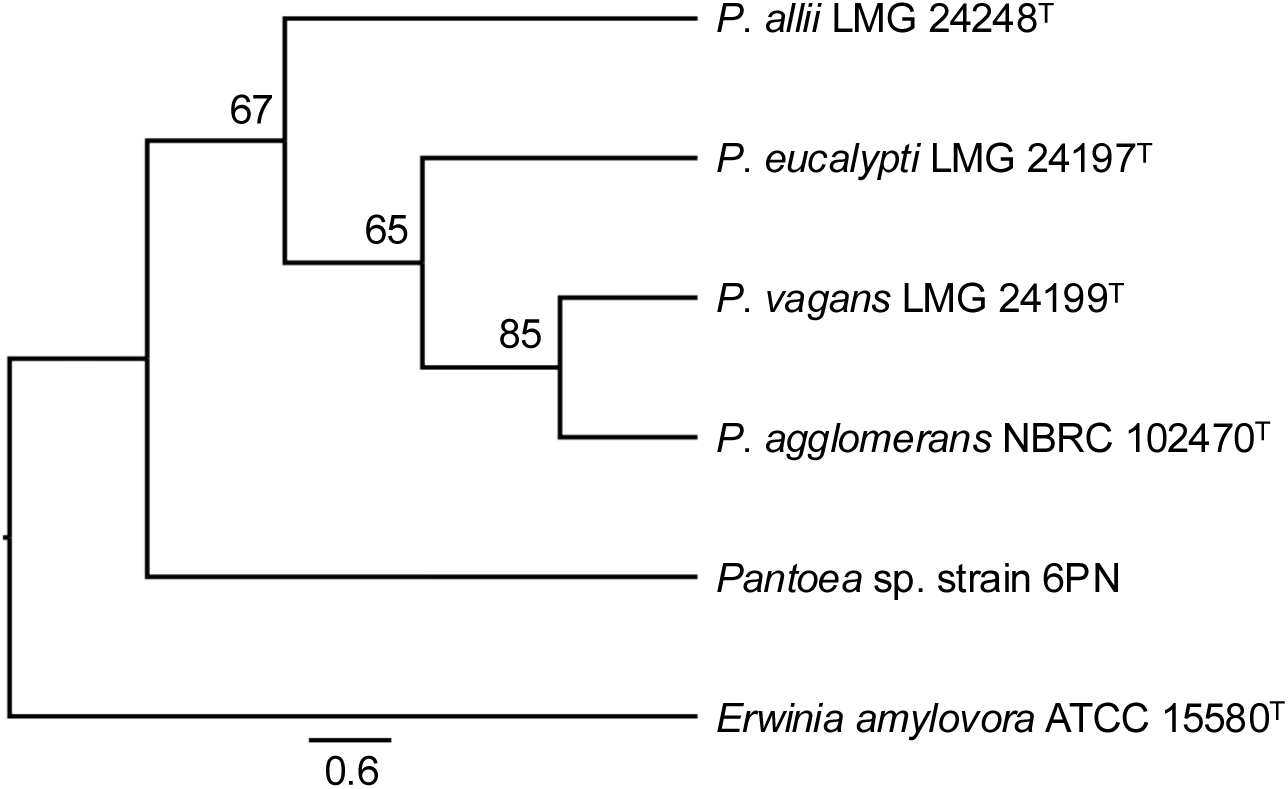
Phylogenetic placement of the quinoa-associated isolate within the genus *Pantoea*. Phylogenetic tree based on 16S rRNA gene sequences showing the relationship between strain 6PN and representative *Pantoea* strains, including type strains. Type strains are indicated by a superscript T. Nucleotide sequences were aligned using MAFFT v7.490 with the L-INS-i algorithm. A poorly aligned terminal region at one end of the alignment was trimmed to standardize the sequence boundaries. The maximum-likelihood tree was constructed using IQ-TREE v3.1.2 with the HKY substitution model selected by the software. Bootstrap values (%) based on 1,000 replicates are shown at branch nodes. *Erwinia amylovora* ATCC 15580ᵀ was used as the outgroup. The scale bar represents 0.6 nucleotide substitutions per site.

**Fig. 2.**
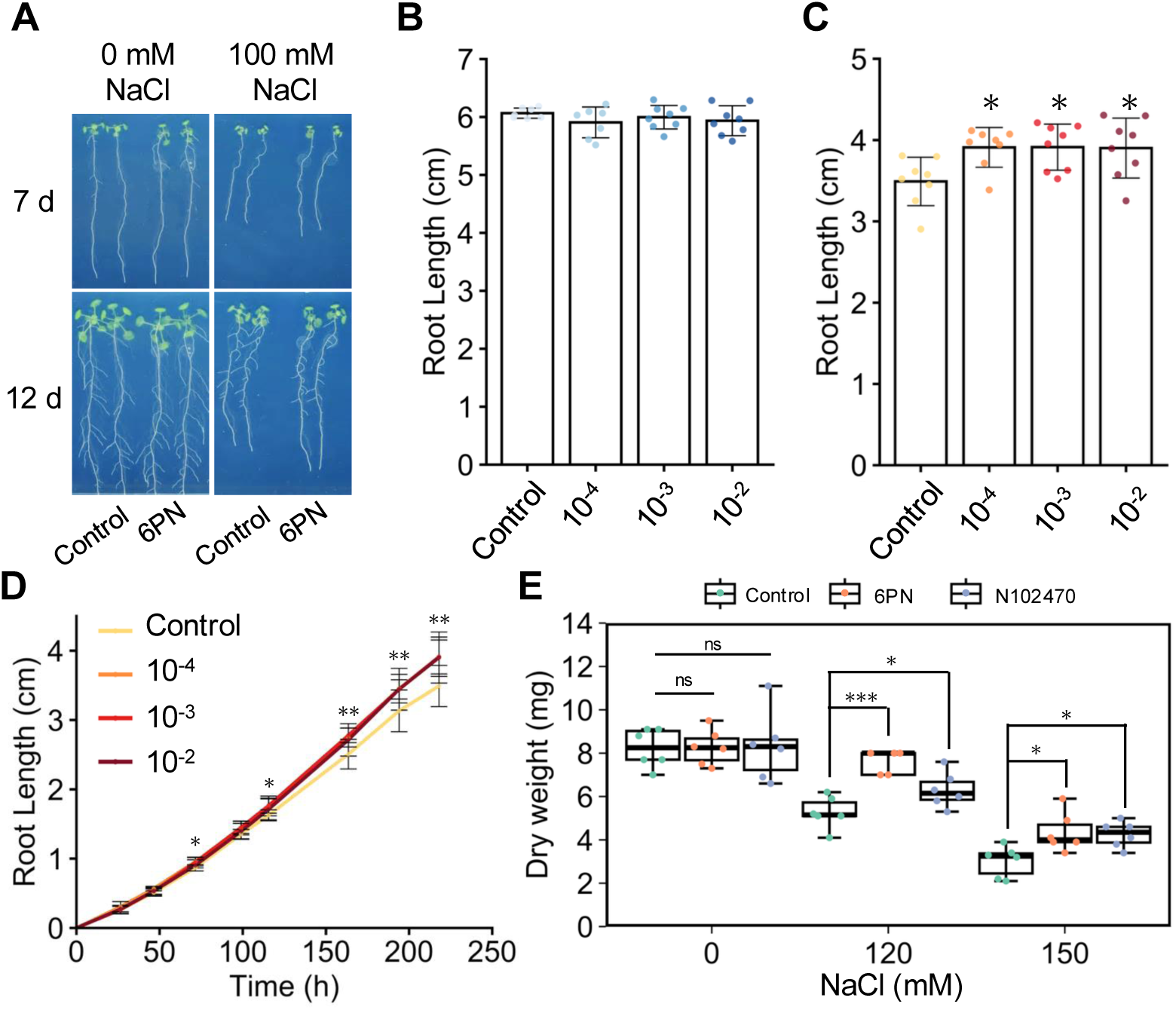
Strain 6PN promotes plant growth specifically under salinity stress conditions. (**A**) Representative images of *Arabidopsis thaliana* seedlings grown under non-saline (0 mM NaCl) or saline (100 mM NaCl) conditions following inoculation with strain 6PN or MgCl₂ buffer as a mock treatment. Seedlings were transferred to treatment medium 4 d after germination and inoculated 2 d later. Images were captured 7 and 12 d after inoculation. (**B**, **C**) Primary root length of *A. thaliana* seedlings measured 7 d after inoculation with different concentrations of strain 6PN under non-saline conditions (**B**) or saline conditions containing 100 mM NaCl (**C**). The x-axis indicates the inoculum concentration of strain 6PN. Data represent means ± SD (*n* = 7–8). Similar results were obtained in three independent experiments. (**D**) Time-course analysis of primary root elongation in *A. thaliana* seedlings grown on medium containing 100 mM NaCl following inoculation with different concentrations of strain 6PN or mock treatment. Primary root length was measured at the indicated time points. Data represent means ± SD (*n* = 8). (**E**) Whole-plant dry weight of *A. thaliana* seedlings inoculated with strain 6PN, *Pantoea agglomerans* NBRC 102470 (N102470), or mock-treated controls under saline conditions. Seedlings were cultivated on medium containing 120 or 150 mM NaCl following bacterial inoculation. Data represent means ± SD (*n* = 5–6). Similar results were obtained in three independent experiments. Asterisks indicate statistically significant differences from the mock-treated control as determined by Dunnett’s multiple-comparison test in panels **B**, **C**, and **E**, or by two-tailed Student’s *t*-test in panel **D** (\**p* < 0.05, \*\**p* < 0.01, \*\*\**p* < 0.001).

When strain 6PN was inoculated onto the root region of *Arabidopsis thaliana* seedlings grown under non-saline conditions, no significant effect on primary root growth was observed compared with mock-inoculated controls during the observation period (Fig. 2A,B). In contrast, under saline conditions containing 100 mM NaCl, inoculation with strain 6PN significantly enhanced primary root elongation at all bacterial concentrations tested (Fig. 2A,C). Time-course analysis further showed that strain 6PN promoted primary root elongation under saline conditions during later stages of seedling growth (Fig. 2D). Consistent with this root growth phenotype, seedlings inoculated with strain 6PN and cultivated on medium containing 120 or 150 mM NaCl exhibited significantly greater whole-plant dry weight than mock-inoculated controls (Fig. 2E). These results indicate that strain 6PN promotes plant growth specifically under salinity stress conditions.

### Strain 6PN exhibits salinity-responsive IAA production

To examine bacterial traits associated with salinity-dependent growth-promoting activity of the quinoa-associated *Pantoea* isolate, we compared its plant growth-promoting traits with those of two closely related reference strains, *Pantoea agglomerans* NBRC 102470 (N102470) and NBRC 12686 (N12686), which showed high 16S rRNA gene sequence similarity to strain 6PN and were available from a public culture collection. We examined phosphate solubilization, carboxymethyl cellulose degradation, siderophore production, and indole-3-acetic acid (IAA) production under both non-saline and saline conditions. No clear differences among the strains were observed for phosphate solubilization, carboxymethyl cellulose degradation, or siderophore production, regardless of the presence or absence of 100 mM NaCl (Fig. 3A–C). In contrast, marked strain-dependent differences were detected in IAA production under saline conditions. Both strain 6PN and strain N102470 showed increased IAA production in the presence of 100 mM NaCl, whereas strain N12686 produced only low levels of IAA under both non-saline and saline conditions (Fig. 3D). Notably, under saline conditions, strain 6PN produced the highest level of IAA among the strains examined, reaching approximately 60 nmol mL⁻¹.

**Fig. 3.**
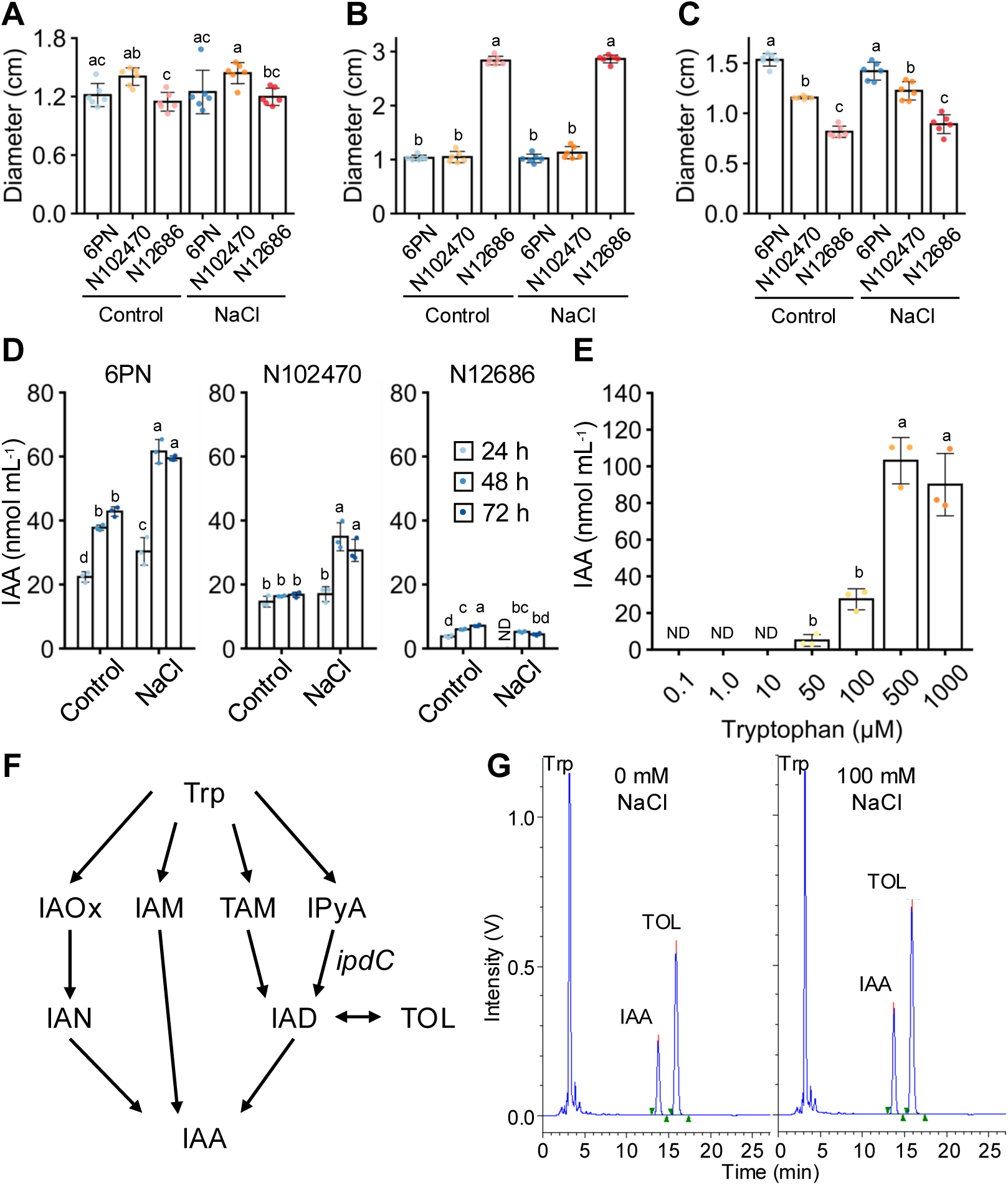
Salinity-responsive auxin production by the quinoa-associated isolate. (**A**) Phosphate solubilization activity of strain 6PN and reference *Pantoea* strains, NBRC 102470 (N102470) and NBRC 12686 (N12686), in the presence or absence of 100 mM NaCl. (**B**) Carboxymethyl cellulose (CMC) degradation activity of strain 6PN and reference strains under non-saline and saline conditions. (**C**) Siderophore production by strain 6PN and reference strains determined using CAS agar medium under non-saline and saline conditions. (**D**) Time-course analysis of indole-3-acetic acid (IAA) production by strain 6PN and reference *Pantoea* strains cultured in the presence or absence of 100 mM NaCl. (**E**) Tryptophan-dependent IAA production by strain 6PN. Cells were cultivated in medium supplemented with increasing concentrations of tryptophan, and IAA accumulation was quantified by HPLC. ND, not detected. (**F**) Putative bacterial IAA biosynthesis pathways and the position of *ipdC* identified in the genome of strain 6PN. Trp, tryptophan; IAOx, indole-3-acetaldoxime; IAM, indole-3-acetamide; TAM, tryptamine; IPyA, indole-3-pyruvic acid; IAN, indole-3-acetonitrile; IAD, indole-3-acetaldehyde; TOL, indole-3-ethanol; *ipdC*, indole-3-pyruvate decarboxylase. (**G**) Representative HPLC chromatograms showing detection of IAA and TOL in culture supernatants of strain 6PN grown in the absence or presence of 100 mM NaCl. Peaks corresponding to tryptophan (Trp), IAA, and TOL were identified using authentic standards. The horizontal and vertical axes indicate retention time (min) and signal intensity (V), respectively. For presentation purposes, only the 0–25 min region of the chromatograms is shown. Under these HPLC conditions, the retention times of Trp, IAA, and TOL were approximately 3.2, 13.7, and 15.9 min, respectively. Data in panels **A**–**C** represent means ± SD (*n* = 6). Data in panels **D** and **E** represent means ± SD (*n* = 3). Different letters indicate significant differences among treatments according to Tukey’s HSD test (*p* < 0.05).

To examine whether salinity-responsive IAA production was accompanied by plant growth-promoting activity under salt stress, we compared the effects of strain 6PN and strain N102470 on the growth of *Arabidopsis thaliana* under saline conditions. Both strains increased whole-plant dry weight under NaCl stress compared with mock-inoculated controls (Fig. 2E). Strain 6PN tended to show a stronger effect than strain N102470 under 120 mM NaCl, whereas the effects of the two strains were similar under 150 mM NaCl. These results show that both strains exhibiting salinity-responsive IAA production promoted plant growth under saline conditions, with strain 6PN showing the highest IAA production among the strains examined.

### Genome analysis identifies putative genes related to auxin production and stress-associated traits in strain 6PN

To further investigate genomic features associated with the physiological properties of strain 6PN, we performed whole-genome sequencing and functional annotation analyses. The assembled genome consisted of a single circular chromosome of 4,142,804 bp and two circular plasmids designated pCq6PN1 and pCq6PN2, with sizes of 322,785 bp and 182,938 bp, respectively. The total genome size was approximately 4.65 Mb, with an average GC content of 54.7% (Fig. 4). Genome assembly completeness was estimated to be 97.46% using the DFAST pipeline. Genome annotation predicted 4,462 putative protein-coding sequences, 22 rRNA genes, 77 tRNA genes, and one transfer-messenger RNA (tmRNA) gene.

**Fig. 4.**
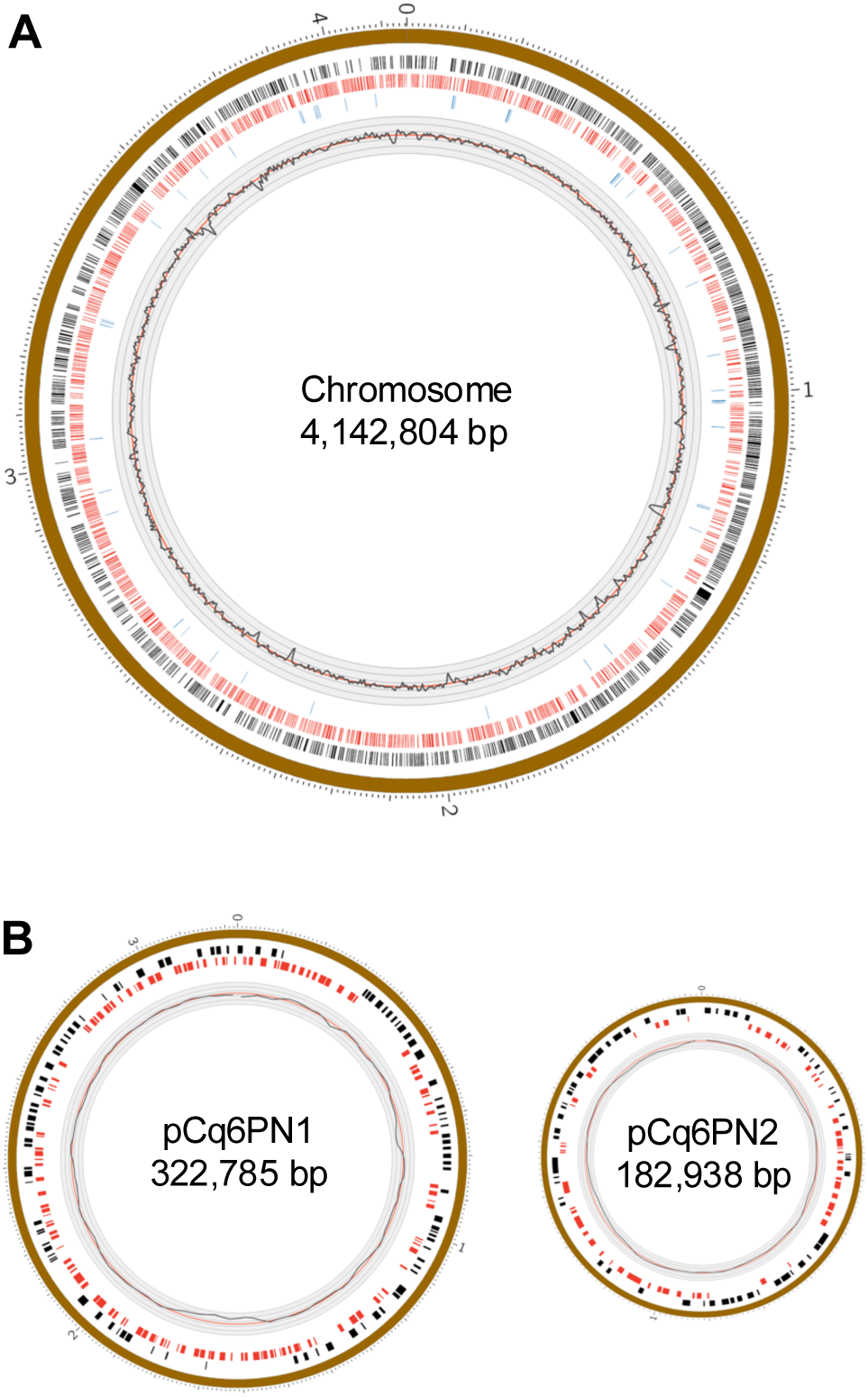
Circular genome maps of strain 6PN. (**A**) Circular map of the chromosome of strain 6PN (4,142,804 bp). From the outermost to innermost circles: genomic coordinates, coding sequences (CDSs) on the forward strand (black), CDSs on the reverse strand (red), tRNA, tmRNA, and rRNA genes (blue), and GC content. GC content was calculated using a 5-kb window with a 2.5-kb step size. Major tick marks indicate 1 Mb intervals. (**B**) Circular maps of plasmid pCq6PN1 (322,785 bp) and plasmid pCq6PN2 (182,938 bp) of strain 6PN. From the outermost to innermost circles: plasmid coordinates, CDSs on the forward strand (black), CDSs on the reverse strand (red), and GC content. Major tick marks indicate 100 kb intervals. Putative genes associated with plant-associated and stress-related functions are summarized in Supplementary Tables 1–6.

Genome annotation identified multiple genes associated with plant-associated and stress-related functions, including putative genes involved in indole-3-acetic acid (IAA) biosynthesis (Supplementary Table 1), phosphorus metabolism (Supplementary Table 2), siderophore production (Supplementary Table 3), reactive oxygen species regulation (Supplementary Table 4), polysaccharide biosynthesis and export (Supplementary Table 5), and flagellar biosynthesis and chemotaxis (Supplementary Table 6). Most of these genes were located on the chromosome. Notably, the genome contained a putative *ipdC* gene encoding indole-3-pyruvate decarboxylase, a key enzyme in the bacterial indole-3-pyruvic acid (IPyA)-associated IAA biosynthesis pathway that catalyzes the conversion of IPyA to indole-3-acetaldehyde (Fig. 3F; Supplementary Table 1). Consistent with the presence of this pathway-related gene, strain 6PN produced IAA in a tryptophan-dependent manner (Fig. 3E). HPLC chromatograms further showed a stronger indole-3-ethanol (TOL)-associated peak in culture supernatants of strain 6PN grown under 100 mM NaCl than under non-saline conditions (Fig. 3G). Because TOL is a metabolite associated with the bacterial IPyA pathway, these observations support the possible involvement of an IPyA-associated bacterial IAA biosynthesis pathway in salinity-responsive IAA production by strain 6PN.

### Phylogenomic analyses indicate that strain 6PN is genomically distinct from representative *Pantoea* species

To further characterize the phylogenomic position of the quinoa-associated isolate, we conducted multilocus sequence analysis (MLSA) and average nucleotide identity (ANI) analysis using representative *Pantoea* strains with available reference genome sequences. MLSA placed strain 6PN within the genus *Pantoea*, in the vicinity of *P. vagans* LMG 24199^T^, *P. agglomerans* NBRC 102470ᵀ, and *P. eucalypti* LMG 24197ᵀ, but on a branch distinct from these representative strains (Fig. 5A). Consistent with this placement, ANI analysis showed that strain 6PN shared approximately 89–91% nucleotide identity with closely related *Pantoea* strains, including 90.8% ANI with *P. vagans* LMG 24199 and 90.5% ANI with *P. agglomerans* NBRC 102470ᵀ (Fig. 5B). These ANI values were substantially lower than the generally accepted species boundary threshold of 95–96%^37^, suggesting that strain 6PN is genomically distinct from the representative *Pantoea* species examined here.

**Fig. 5.**
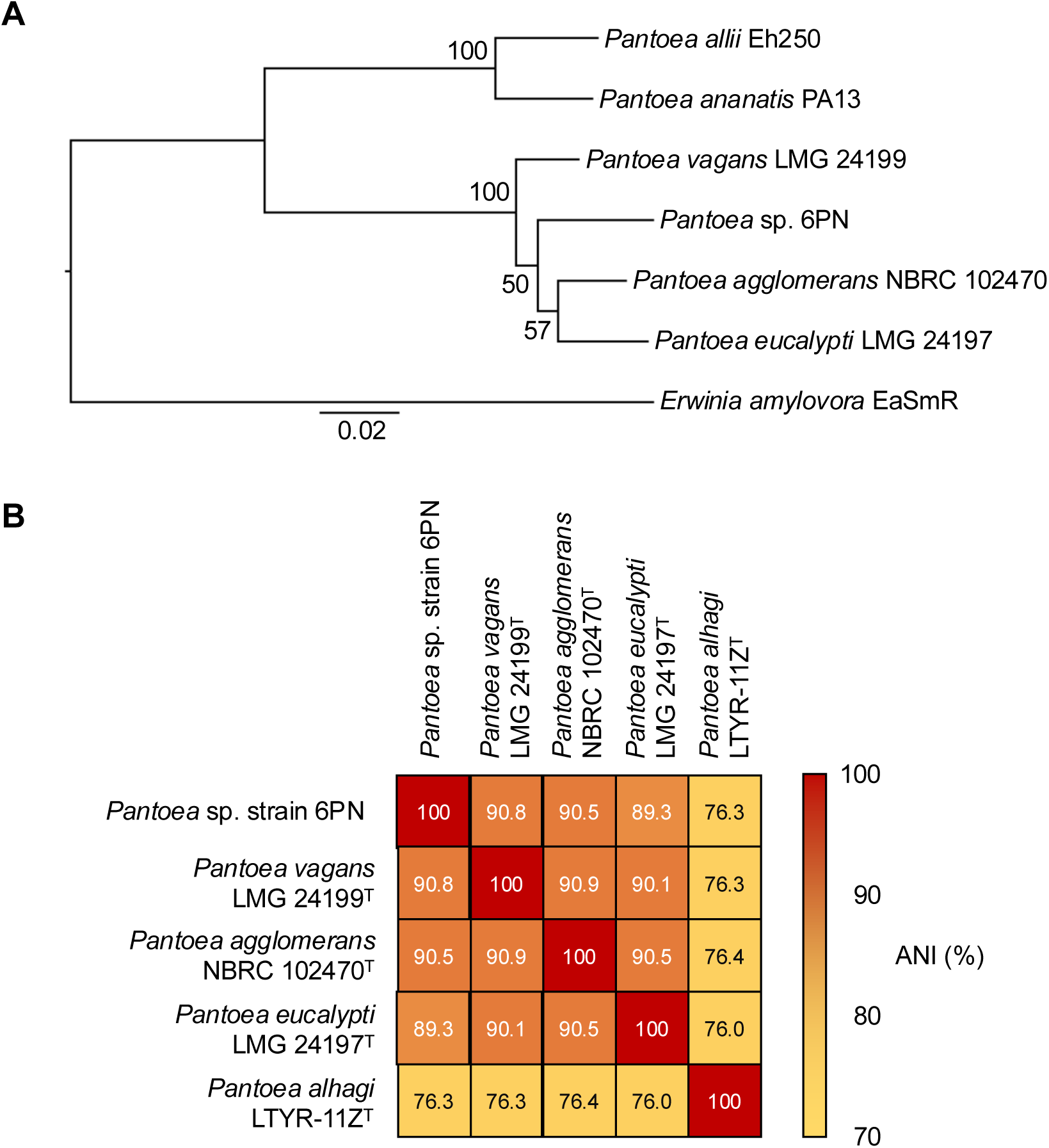
Phylogenomic analyses show that strain 6PN is genomically distinct from representative *Pantoea* species. (**A**) Multilocus sequence analysis (MLSA) showing the phylogenomic position of strain 6PN relative to representative *Pantoea* strains with reference genome sequences. The maximum-likelihood tree was constructed from concatenated coding sequences of six housekeeping genes (*rpoD*, *rpoB*, *recA*, *dnaN*, *dnaA*, and *atpD*) using IQ-TREE v3.1.2 with the TIM2+F+I+G4 model selected by the software. Bootstrap values (%) based on 1,000 replicates are shown at branch nodes. *Erwinia amylovora* EaSmR was used as the outgroup. The scale bar represents nucleotide substitutions per site. (**B**) Pairwise average nucleotide identity (ANI) values between strain 6PN and representative *Pantoea* strains calculated using OrthoANI v0.5.0. Strains shown in the ANI matrix were selected based on their close phylogenetic proximity to strain 6PN and the availability of whole-genome assemblies. Colors indicate ANI values according to the scale shown on the right. ANI values below the generally accepted species boundary threshold of 95–96% indicate that strain 6PN is genomically distinct from the representative *Pantoea* species examined here.

### Strain 6PN associates with spatially separated plant tissues under saline conditions

To evaluate the ability of strain 6PN to associate with spatially separated plant tissues under saline conditions, we used an *Arabidopsis thaliana* trench-plate assay in which GFP-labeled bacterial cells were inoculated onto either the leaf or root region of seedlings grown on 0.5× MS medium supplemented with 120 mM NaCl (Fig. 6A). Four to five days after inoculation onto leaf tissues, GFP fluorescence derived from strain 6PN-GFP was detected on root surfaces, despite the physical separation of leaf and root regions by the trench structure (Fig. 6B). Similar GFP fluorescence signals were also observed for the reference strain *Pantoea agglomerans* NBRC 102470-GFP. These observations indicate that strain 6PN can be detected on root surfaces following inoculation onto aerial plant tissues in this assay system.

**Fig. 6.**
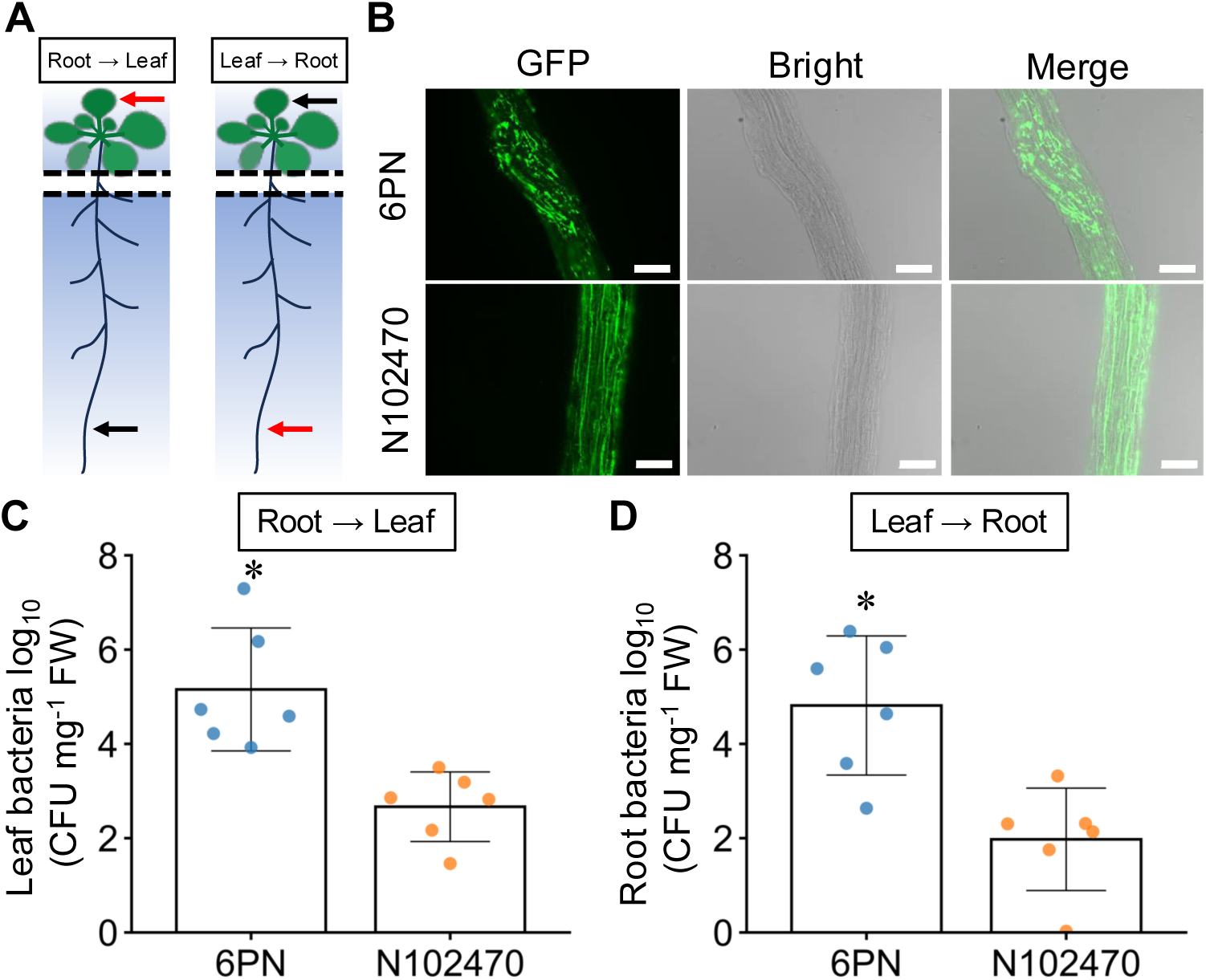
Higher recovery of strain 6PN from spatially separated plant tissues under saline conditions. (**A**) Schematic illustration of the trench-plate assay used to evaluate bacterial association with spatially separated leaf and root tissues under saline conditions. Black and red arrows indicate the inoculation site and the tissue used for bacterial detection, respectively. (**B**) Representative GFP fluorescence images showing bacterial association with root surfaces by strain 6PN-GFP and *Pantoea agglomerans* NBRC 102470-GFP following inoculation onto leaf tissues. GFP fluorescence, bright-field, and merged images are shown. Scale bars, 100 μm. (**C**) Quantification of bacteria recovered from leaf tissues following inoculation onto root tissues (Root → Leaf). Bacterial abundance was determined as GFP-labeled CFUs on selective medium containing gentamicin, normalized to tissue fresh weight, and expressed as log₁₀ CFU mg⁻¹ fresh weight. (**D**) Quantification of bacteria recovered from root tissues following inoculation onto leaf tissues (Leaf → Root). Bacterial abundance was determined as GFP-labeled CFUs on selective medium containing gentamicin, normalized to tissue fresh weight, and expressed as log₁₀ CFU mg⁻¹ fresh weight. Data in panels **C** and **D** represent means ± SD (*n* = 6). Asterisks indicate significant differences between strain 6PN and NBRC 102470 as determined by two-tailed Student’s *t*-test (*p* < 0.05).

To quantitatively assess bacterial recovery from spatially separated tissues, leaf or root tissues were excised 5–7 days after inoculation, homogenized, and plated onto selective medium for colony counting. Following inoculation onto root tissues, both strain 6PN-GFP and strain NBRC 102470-GFP were recovered from leaf tissues (Fig. 6C). Conversely, when bacteria were inoculated onto leaf tissues, both strains were recovered from root tissues (Fig. 6D). In both experimental systems, significantly higher numbers of colony-forming units (CFUs) were recovered from plants inoculated with strain 6PN-GFP than from those inoculated with strain NBRC 102470-GFP. These results suggest that strain 6PN is recovered from spatially separated plant tissues at higher levels than NBRC 102470 under saline conditions in the *Arabidopsis* trench-plate assay.

## Discussion

Here, we identified strain 6PN, a quinoa-associated *Pantoea* isolate obtained from surface-sterilized seedlings derived from a long-term laboratory-propagated quinoa line, as an isolate that promoted the growth of *Arabidopsis thaliana* under saline conditions (Figs. 1–2). A central finding of this study is that IAA production by strain 6PN increased under saline conditions (Fig. 3). Bacterial indole-3-acetic acid (IAA) production is a well-known plant growth-promoting trait^9,38^, but it is often evaluated under standard culture conditions. In the present study, strain 6PN showed increased IAA production in the presence of NaCl, and strain NBRC 102470 showed a similar but weaker response, whereas strain NBRC 12686 produced only low levels of IAA under the conditions tested (Fig. 3D). Together, these results indicate that IAA production varies among *Pantoea* strains and is enhanced by salinity in some strains. The putative *ipdC* gene identified in strain 6PN (Fig. 3F; Supplementary Table 1) and the stronger TOL-associated peak under saline conditions (Fig. 3G) support the possible involvement of an IPyA-associated pathway in IAA production by strain 6PN, but this remains to be functionally tested^9,38^. These results identify salinity-responsive IAA production as a prominent physiological feature of strain 6PN.

The growth-promotion data are consistent with the salinity-responsive IAA production observed in strain 6PN. Strain 6PN promoted primary root elongation and whole-plant dry weight under saline conditions, whereas no significant growth-promoting effect was observed under non-saline conditions (Fig. 2). In addition, strain NBRC 102470 also increased whole-plant dry weight under salt stress, although it produced less IAA than strain 6PN (Figs. 2, 3D). These findings suggest that salinity-responsive IAA production is a prominent bacterial trait associated with growth-promoting activity under salt stress, while also indicating that additional bacterial or plant factors may contribute to the observed growth response.

Genome analysis provides additional context for the physiological properties of strain 6PN. In addition to the putative *ipdC* gene, strain 6PN carried genes related to stress responses, nutrient acquisition, polysaccharide biosynthesis and export, flagellar biosynthesis, and chemotaxis (Fig. 4; Supplementary Tables 1–6). These annotations provide genomic candidates for traits that may support bacterial survival, stress responsiveness, or plant association under saline conditions. Phylogenomic analysis and ANI comparisons further indicated that strain 6PN is genomically distinct from the representative *Pantoea* species examined here (Fig. 5), although formal taxonomic assignment will require additional characterization^37^.

The *Arabidopsis* trench-plate assay showed that strain 6PN-GFP was recovered from spatially separated plant tissues at higher levels than strain NBRC 102470-GFP under saline conditions (Fig. 6). This higher recovery suggests that strain 6PN is more effectively retained by, or distributed across, plant tissues in this assay system. The presence of genes related to flagellar biosynthesis and chemotaxis in the strain 6PN genome is consistent with possible plant-associated behavior, although the present assay does not resolve the precise route of bacterial spread. Together, these results show that strain 6PN is recovered at higher levels than the reference strain from spatially separated plant tissues under saline conditions.

Plant-associated microorganisms have been implicated in plant performance under stressful environments, including saline or otherwise harsh habitats^5,20,21^. Quinoa provides a useful source material for exploring plant-associated bacteria because it has strong intrinsic salinity tolerance, including in lines maintained under long-term laboratory propagation^17,18,40^. In this study, strain 6PN was isolated from surface-sterilized seedlings derived from a long-term laboratory-propagated quinoa line. This result supports the use of long-term laboratory-propagated quinoa as a source for isolating bacteria with stress-responsive physiological traits, while the relationship between strain 6PN and quinoa salinity tolerance remains to be examined. Because preliminary quinoa seedling assays did not provide a suitable quantitative readout for bacterial growth promotion under salt-stress conditions, the use of *Arabidopsis* as a model system allowed evaluation of strain 6PN activity under defined saline conditions. Taken together, this study shows that a quinoa-associated *Pantoea* isolate can exhibit salinity-responsive IAA production together with plant growth-promoting activity under salt stress in a model plant system. These findings highlight salinity-responsive bacterial physiology as a useful perspective for exploring plant-associated bacteria under salt-stress conditions.

## Data availability statement

The genome sequence data generated in this study have been deposited in DDBJ under BioProject number PRJDB42378 (AP048863-AP048865).

## Acknowledgements

We thank Dr. Jennifer L. Morrell-Falvey (Oak Ridge National Laboratory, USA) for kindly providing the plasmid pBBR1-MCS5-GFP. We thank the staff of JIRCAS, M. Toyoshima, Y. Nakamura, J. Baba, M. Ikegami, I. Gejima, Y. Nonoue, Y. Takiguchi, A. Aoyama, T. Nada, M. Karasawa, N. Saito, N. Ohmiya, Y. Masamura, K. Ozawa, Y. Shirai, N. Hisatomi, W. Kawakami, K. Shimizu, A. Sugitani, and Y. Kida for their excellent technical assistance.

## Author contributions

YM: Conceptualization, Data curation, Formal analysis, Investigation, Methodology, Visualization, Writing – original draft, Writing – review & editing. TK: Data curation, Formal analysis, Investigation, Methodology, Validation, Visualization, Writing – original draft, Writing – review & editing. HD: Investigation, Data curation, Formal analysis, Visualization, Writing – original draft, Writing – review & editing. YK: Formal analysis, Investigation, Resources, Data curation, Visualization, Validation, Methodology, Writing – original draft, Writing – review & editing. YF: Conceptualization, Methodology, Funding acquisition, Project administration, Resources, Supervision, Validation, Formal analysis, Visualization, Writing – original draft, Writing – review & editing.

## Additional information Supplementary Information

**Supplementary Table 1**

Identification of the putative ipdC gene in strain 6PN.

**Supplementary Table 2**

Putative genes associated with phosphate solubilization and transport in strain 6PN.

**Supplementary Table 3**

Putative genes associated with siderophore biosynthesis and ferric iron transport in strain 6PN.

**Supplementary Table 4**

Putative genes associated with reactive oxygen species (ROS) regulation in strain 6PN.

**Supplementary Table 5**

Putative genes associated with polysaccharide biosynthesis and export in strain 6PN.

**Supplementary Table 6**

Putative genes associated with flagellar biosynthesis and chemotaxis in strain 6PN.

**Correspondence** and requests for materials should be addressed to Y.F.

